# Investigating apparent differences between standard DKI and axisymmetric DKI and its consequences for biophysical parameter estimates

**DOI:** 10.1101/2023.06.21.545891

**Authors:** Jan Malte Oeschger, Karsten Tabelow, Siawoosh Mohammadi

## Abstract

**Purpose:** Identify differences between the acquisition-time efficient axisymmetric diffusion kurtosis imaging (DKI) model and standard DKI and their consequences on biophysical parameter estimates using standard DKI parameters as the ground truth.

**Methods:** Noise-free, synthetic diffusion MRI (dMRI) human brain data are generated using standard DKI and fitted with axisymmetric DKI and standard DKI. Then, the five axisymmetric DKI tensor metrics (AxTM), the parallel and perpendicular diffusivity and kurtosis and mean of the kurtosis tensor, attainable with both DKI models are computed. Next, the five biophysical parameters axon water fraction and dispersion, extra axonal parallel and perpendicular diffusivity and intra axonal parallel diffusivity are estimated from the AxTM using the WMTI-Watson model. Finally, the number of substantially differing voxels (SDV), defined as voxels where estimation results of both DKI models differ more than 5%, is calculated for the AxTM and the biophysical parameters.

**Results:** For the AxTM, the number of SDV was biggest for the parallel (26%) and perpendicular (51%) kurtosis while the other three AxTM had very few SDV (less than 5%). The biophysical parameters had much more SDV than the AxTM from which they were computed, ranging from 29% to 50%.

**Conclusion:** Axisymmetric DKI is a viable alternative to standard DKI in studies focusing on effects based on the parallel and perpendicular diffusion and mean of the kurtosis tensor. However, our findings urge caution when using axisymmetric DKI to investigate effects based on the parallel and perpendicular kurtosis or use it to estimate the biophysical parameters.

## 1 Introduction

Diffusion kurtosis imaging (DKI) has increasingly been used to study the neuronal tissue microstructure and derive biophysical parameters relevant for understanding brain function and impact of disease ^1;2;3^ since ten years. DKI is a more complex extension to the well known diffusion tensor imaging (DTI) framework and provides diffusion kurtosis metrics that are also sensitive to the tissue microstructure and can provide complementary information ^4;5;6^ to DTI. However, the increased complexity goes hand in hand with an increase in acquisition time. Since time is a limited resource in scientific and especially clinical settings, needing more time poses a major hurdle for a more extensive implementation and application of DKI instead of DTI.

Axisymmetric DKI was recently introduced as a more acquisition-time efficient DKI model^7;8;9^ because the reduced parameter space of axisymmetric DKI (8 parameters) can be fitted with less data than is needed for fitting the whole standard DKI parameter space (21 parameters). Axisymmetric DKI reduces the parameter space by imposing additional symmetry assumptions, i.e., axisymmetrically organized fibers in the imaged tissue structure. However, violation of these additional symmetry assumptions might lead to a deviation of axisymmetric DKI fit results from their standard DKI reference counterpart. An important subset of the parameters that exist in both DKI frameworks are the five axisymmetric DKI tensor metrics (AxTM), the parallel and perpendicular diffusivity (*D*_‖_ and *D*_⊥_) and kurtosis (*W*_‖_ and *W*_⊥_) and mean of the kurtosis tensor 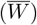. The five AxTM are also directly related to the five parameters of the biophysical standard model ^10^ axon water fraction f, axon dispersion *κ*, parallel and perpendicular extra-axonal diffusivities *D*_*e*,‖_ and *D*_*e*,⊥_ and intra axonal diffusivity *D*_a_ here estimated with the WMTI-Watson model ^11;12^. The AxTM are ideally suited to assess potential differences between fit results estimated with axisymmetric DKI and standard DKI because they are frequently used invariants that can be estimated by both models. Furthermore, they are used to compute the biophysical parameters and therefore allow to study propagation of the apparent differences in the AxTM into them.

Our study seeks to explore the differences of parameter estimation results of both DKI models and its propagation into the biophysical parameters based on synthetic, noise-free, healthy human, in-vivo dMRI data. To this end, standard DKI is used as a forward model for generating the synthetic dMRI data to explicitly include complex, non-axisymmetric fiber configurations. It has been suggested that the symmetry assumptions made in axisymmetric DKI are likely a reasonable approximation to diffusion in major white matter fiber bundles ^7^ which is why the focus of this study is white matter. However, even in white matter voxels, the symmetry assumption in axisymmetric DKI might be violated, e.g., in areas where fiber crossings occur ^13^. This can lead to deviations between parameters estimated with axisymmetric DKI and standard DKI. A hypothesis investigated here is that any observed, apparanet deviation between both DKI model variants is due to an error in axisymmetric DKI rooted in an underlying complex and thus non-axisymmetric fiber configuration. To this end a fractional anisotropy (FA) threshold is used to identify voxels with highly anisotropic diffusion, i.e., less complex diffusion patterns following the findings in ^14^ and thus a higher probability of fulfilling the assumption of an axisymmetric fiber configuration. Throughout the study, the estimates of the five AxTM or the biophysical parameters based on standard DKI are used as a ground truth reference and compared to the results found by axisymmetric DKI on a voxel-wise basis.

## 2 Methods

A detailed description of the standard DKI model and the axisymmetric DKI model is provided in the Supporting Information Section S1.1 and Section S1.2 but can also be found in ^7^ and ^15^.

### 2.1 Dataset

#### Acquisition

Multi-shell, in-vivo dMRI data with 153 diffusion gradient directions and b-values of 0, 550, 1100 and 2500 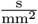 were acquired from a healthy volunteer at 3T with: FOV of 200×203×170mm^3^ at 1.7mm isotropic resolution and 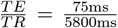. The following pre-processing was applied to the dMRI data (in that order): eddy current and motion artifact correction, susceptibility artifact correction and Rician bias correction acting on the signals of the individual diffusion weighted images ^16^. Additionally, we acquired multi parameter mapping data^17^ on the same subject and calculated the R1 map using the hMRI toolbox ^18^.

#### Generation of synthetic data

The acquired multi-shell dMRI data were fitted with standard DKI to obtain the 22 standard DKI tensor metrics which were then used for generation of noise-free, synthetic dMRI data using standard DKI as a forward model.

### 2.2 Biophysical parameters

The framework presented in ^10^, here refereed to as “WMTI-Watson”, was used to establish an analytical connection between the five AxTM Ω = {*D*_‖_, *D*_⊥_, *W*_‖_, *W*_⊥_, 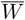} and the biophysical parameters β = {AWF, κ, *D*_e,⊥_, *D*_e,‖_, *D*_*a*_}.

First κ is estimated by minimizing objective function Eq. (S1.13) (Supporting Information). It was feasible to optimize this problem over a discrete, linearly sampled range of [0 ≤ κ ≤ 50] because it depends on one non-negative parameter κ. This procedure was faster and more precise compared to using the available MATLAB solvers. Using κ, the other AxTM can be computed following the formulas in Section S1.3, Supporting Information. There are at least two solutions^10^ (“branches”) to the optimization problem, but here only the results of the branch labeled “+” is reported. This branch choice corresponds to assuming 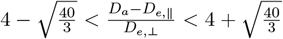 associated with *D*_*a*_ > *D*_‖_ and also labeled η = 1 in ^19^ or ζ = + in ^10^, see Section 4.2 for a further discussion.

### 2.3 Computation of difference between both DKI models and substantially differing voxels (SDV)

The estimated parameters, either the set of AxTM Ω or the set of biophysical parameters β were estimated based on standard DKI and axisymmetric DKI and compared using the voxel-wise absolute percentage error (A-PE):

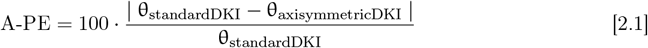

Here θ is an element of either Ω or β, the subscript indicates whether the parameter was estimated based on standard DKI or axisymmetric DKI. If A-PE > 5%, the corresponding voxel was classified as a “substantially differing voxel” (SDV). The study focused on white matter only, to obtain the white matter mask, we segmented the R1 map into tissue probability maps (TPM) and thresholded the white matter TPM (TPM > 0.9), see green contour of Figure 1. To summarize the results we estimated a) the number of SDV in percent and b) the median difference in the population of SDV. We implemented the condition θ ∈ Ω ≥ 0 and θ ∈ β ≥ 0 because diffusivity values smaller 0 are non-physical and AWF and κ are ≥ 0 by definition. Furthermore, kurtosis estimates in the healthy brain have been found ^20^ well above 0.

**Figure 1.**
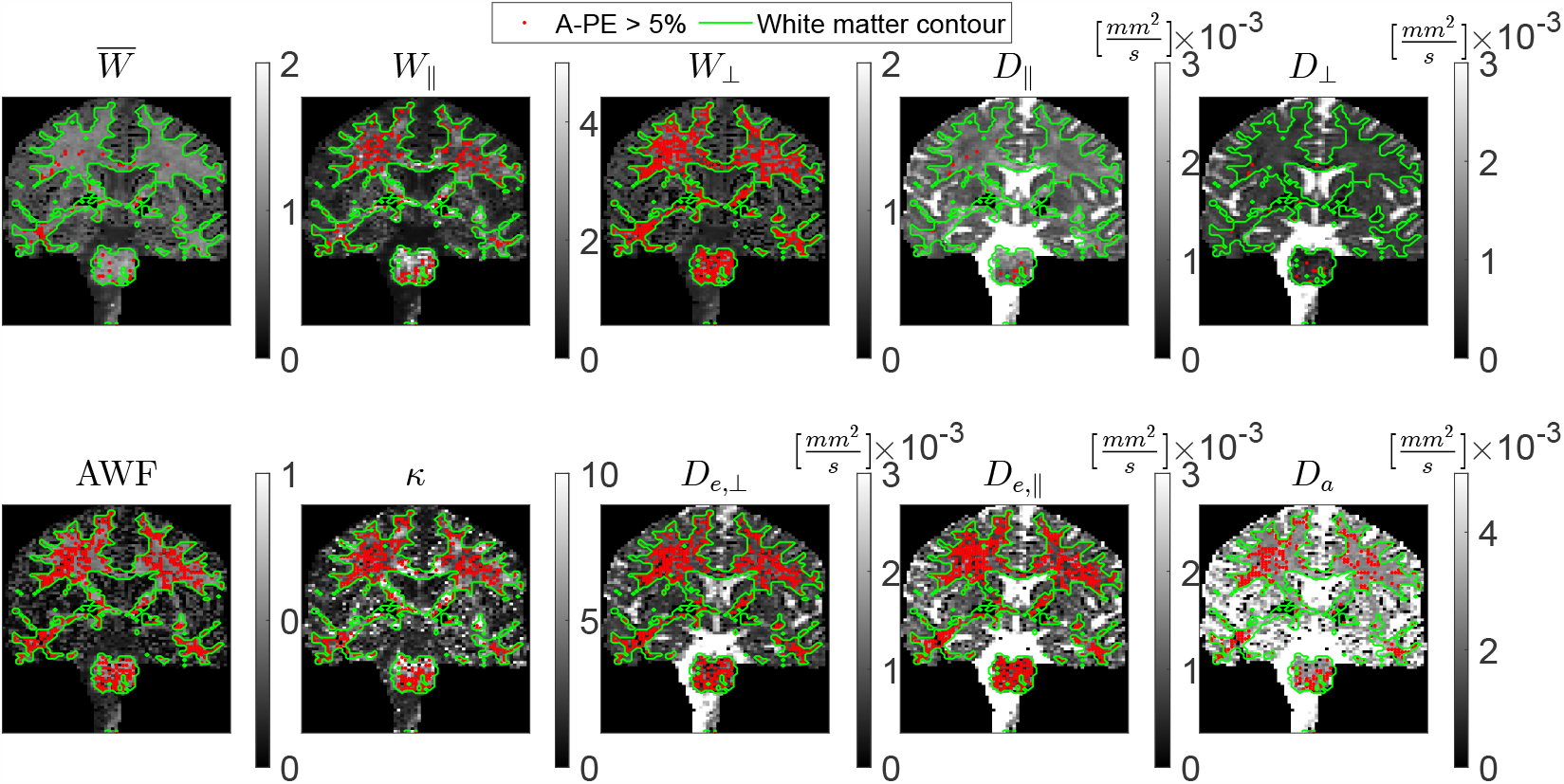
Examples of the AxTM (top) and biophysical parameters (bottom) in a slice of the human brain dMRI data used in this study. The green contour outlines the white matter contour, the red dots indicate voxels where the A-PE≥ 5% (“substantially differing voxels”). The red barplots of the top row of Figure 2 show the percentage of substantially differing voxels in the whole white matter.

#### Influence of FA threshold on number of substantially differing voxels

Either the whole white matter or the whole white matter masked with a fractional anisotropy (FA) mask FA≥ 0.55 was investigated. The FA mask was introduced to investigate the effects of the fiber complexity on our results, the threshold of 0.55 was based on the FA of an unidirectional phantom, see ^14^, its function is to guarantee the existence of a well defined main diffusion pathway and therefore a less complex diffusion pattern where axisymmetric DKI’s assumptions are likely fulfilled.

#### Inter-dependence of A-PE and difference in main fiber orientation

An earlier work ^21^ has analytically shown that axisymmetric DKI and standard DKI should produce the same results if two pre-conditions are fulfilled: a) the log of the signals is being fitted and b) the axis of symmetry 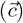 and the first eigenvector 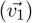, two measures for the main fiber orientation in both DKI models, are identical. To investigate a possible inter-dependency between the A-PE and the difference between 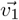 and 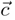, the angle *ϕ* between 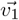 and 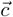 was calculated according to: 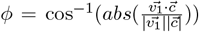 and plotted against the A-PE as a scatter density plot for each parameter.

## 3 Results

### 3.1 Summary measures: Number of substantially differing voxels highly parameter dependent and biophysical parameters affected the most. Median A-PE similar across all parameters

#### Differences between AxTM across the white matter using the two DKI models

Figure 1 shows the spatial distribution of substantially differing voxels (SDV, see Section 2.3) as red voxels in a slice of the five AxTM and biophysical parameters, Figure 2 summarizes the number of SDV and the median A-PE in that population using barplots.

**Figure 2.**
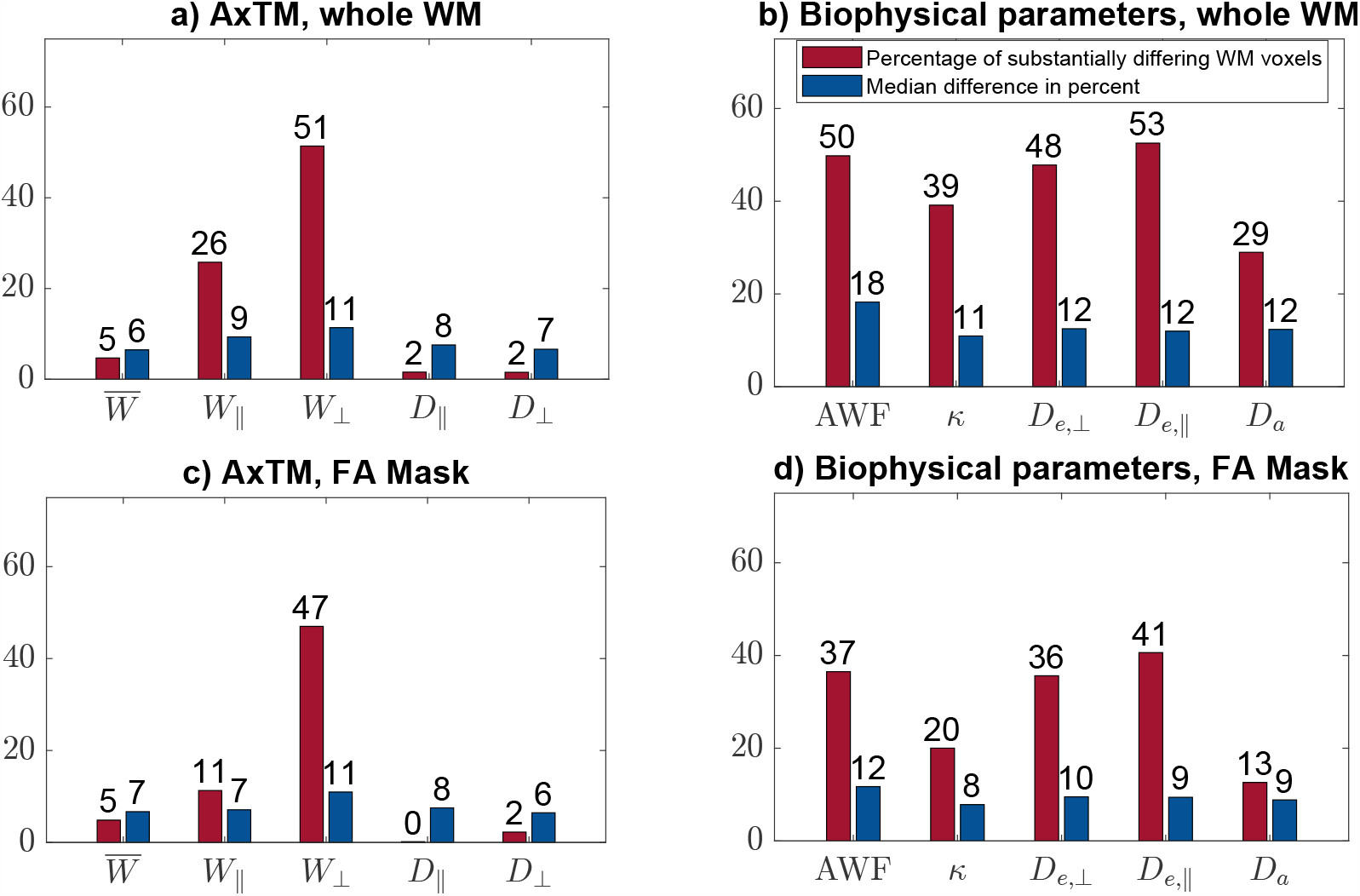
Summarizing barplots for the AxTM (left) and biophysical parameters (right). Shown are the number of substantially differing voxels (red barplots) and the median difference in those voxels (blue barplots). Top row shows the analysis in the whole white matter, the bottom row shows the results in white matter with an applied FA mask (FA ≥ 0.55).

The number of SDV in the white matter voxels is highly parameter dependent (Figure 2), e.g., only 2% for *D*_‖_, *D*_⊥_ and 5% for the mean of the kurtosis tensor 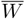. The *W*_‖_ (26%) and *W*_⊥_ (51%) were affected much more, see also spatial distribution of SDV (red voxels in Figure 1). However, the median difference of the SDV across the AxTM was more similar and ranged between 6% 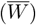 and 11% (*W*_⊥_).

#### Differences between biophysical parameters across the white matter based on the two DKI models

Both the number of SDV and the median A-PE in the SDV population (11% to 18%) was higher for the biophysical parameters than the AxTM. Again, the percentage of SDV was parameter dependent and spanned from 29% (*D*_*a*_) to 53% (*D*_*e,‖*_), see Figure 2b). *D*_*e,‖*_, *D*_*e,⊥*_ and AWF had most SDV while *D*_*a*_ and κ had the least. Figure S2, Supporting Information Section S1.5 documents the underlying A-PE histograms distributions.

For neither the AxTM nor the biophysical parameters Figure 1 revealed a spatial distribution pattern of the SDV in the depicted white matters slice.

### 3.2 Influence of FA threshold on number of substantially differing voxels

#### AxTM

Introduction of an FA threshold reduced the number of SDV for all AxTM, especially for *W*_‖_ while the spread of the median A-PE remained the same, between 6% to 11%, see Figure 2c). The number of *W*_⊥_ SDV only changed slightly (from 51% to 47%) when introducing the FA threshold.

#### Biophysical parameters

The FA threshold reduced the number of SDV in all parameters significantly and the spread of the median A-PE was reduced from (11% to 18%) to (8% - 12%), see Figure 2d).

### 3.3 Inter-dependence of A-PE and difference in main fiber orientation

In almost all AxTM voxels the angle *ϕ* was greater 0, see x-axes of Figure S3 (Supporting Information), indicating that the main fiber orientations estimated by both DKI models were almost never identical. This violates one of the necessary presumptions named in ^21^. The scatter density plots in Figure S3 indicate that the kurtosis metrics *W*_⊥_ and *W*_‖_ had the highest inter-dependency between A-PE and *ϕ*.

## 4 Discussion

This work demonstrated that the deviation between axisymmetric DKI and standard DKI was not the same for all axisymmetric DKI tensor metrics (AxTM). For *D*_‖_, *D*_⊥_ and 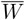 we found little differences between the two DKI models, while *W*_⊥_ and *W*_‖_ showed larger differences. All five axisymmetric DKI based biophysical parameters were strongly different from their standard DKI based counterparts. Introduction of an FA threshold was able to mitigate the observed differences between both DKI models (especially for *W*_‖_ and the biophysical parameters) suggesting that, fiber complexity might be one cause of the observed differences between both DKI models.

### 4.1 Differences between AxTM across the white matter using the two DKI models

The AxTM capture different properties of diffusion in tissue and it is not surprising that the observed differences between both DKI models is AxTM dependent. We used the number of substantially differing voxels (SDV) to quantify the differences between axisymmetric DKI and standard DKI. We found that the diffusion parameters *D*_⊥_ and *D*_‖_ and the mean of the kurtosis tensor 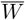 have very few SDV compared to *W*_⊥_ and *W*_‖_.

With a median A-PE of 7% to 11% the error made in axisymmetric DKI might be acceptable depending on the application. Purely judging from the number of SDV, the diffusion parameters were “safest” with only 2% of SDV, followed by the mean of the kurtosis tensor 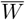 with 5%. It can generally be expected that the kurtosis parameters are more sensitive to a model error since they are quadratic in the b-value b compared to the linear diffusion parameter counterparts. The propagation of error when fitting the axisymmetric DKI model to dMRI data therefore will be more severe for the kurtosis parameters. Interestingly, the number of SDV quintuples from 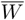 to *W*_‖_ and doubles from *W*_‖_ to *W*_⊥_. The reason for this trend still needs to be explored.

### 4.2 Differences between biophysical parameters across the white matter based on the two DKI models

The number of SDV were significantly enhanced in the biophysical parameters (in some cases up to 53%, see Section 4.3) compared to the AxTM from which they were computed. A reason for the enhancement might be that all five AxTM are required to estimate the biophysical parameters, see Section S1.3 (Supporting Information), and the connection is complex and non-linear. The observed AxTM differences could therefore be amplified due to non-linear effects but also synergistically enhance the number of SDV in the biophysical parameters. However, the found median A-PE of 11% to 18% might be acceptable depending on the study.

Also, similar to ^19^, here the “−” branch tended to yield physically unfeasible, constant high (50) κ values where the objective function did not have a well defined minimum for κ. Furthermore, for this branch *D*_e,‖_ > *D*_*a*_ which in healthy white matter was found to be the biologically invalid solution by most studies^22^. We therefore did not report the results of the − branch.

### 4.3 Influence of FA threshold on number of substantially differing voxels

The difference between standard DKI and axisymmetric DKI may be linked to fiber complexity that brakes the symmetry assumptions of axisymmetric DKI. A proxy for the fiber complexity is the fractional anisotropy (FA) that can be used to identify highly anisotropic diffusion. This was, for example, demonstrated for a unidirectional phantom ^14^.

The FA threshold generally reduced the number of SDV in both the AxTM and the biophysical parameters, supporting the hypothesis that differences between both DKI models are linked to the underlying fiber complexity. However, the FA threshold effectiveness was parameter dependent and, e.g., worked particularly well for *W*_‖_ where it more than halved the number of SDV while it had only small effects on *W*_⊥_. The smaller effectiveness on *W*_⊥_ could come from axisymmetric DKI oversimplifying estimation of *W*_⊥_ since it is directly estimated as a model parameter instead of calculated from three separate tensor metrics as in standard DKI^23^. Note that in-vivo tissue FA can also be influenced by other factors like the degree of myelination or axon density and radius. This means that the conclusion “if FA ≥ 0.55 then the voxel has a unidirectional fiber configuration” is not necessarily strictly true and voxels above the FA threshold might still have a complex fiber structure.

### 4.4 Inter-dependence of A-PE and difference in main fiber orientation

It was shown that axisymmetric DKI produces the same results as standard DKI if two requirements^21^ are met, see Section 2.3. Fulfillment of condition b) was not explicitly checked in^21^ where differences between axisymmetric DKI and standard DKI were reported. The degree to which the main fiber orientations, estimated with both DKI models, differ can be quantified with the angle ϕ between them. The majority of angles ϕ were between ≈ 1 to 5 degrees in white matter (Figure S3, Supporting Information) demonstrating that condition b) is not fulfilled in most cases.

Investigating the dependency of the A-PE on angle *ϕ* using density scatter plots showed an inter-dependency predominantly for *W*_⊥_ and *W*_‖_, Figure S3 (Supporting Information). For these parameters these findings indicate that at least to some extent, there is a causal relationship between *ϕ* and the A-PE.

It could be ruled out that the observed differences between both DKI models in this study is only due to violating condition a) of ^21^ by implementing a log-of-signals fit demonstrating that this fit implementation still produced different fit results for both DKI models, see Supporting Information Section S1.4.

## 5 Conclusion

Axisymmetric DKI offers advantages like a reduced data demand and noise robustness that are relevant in scientific and clinical practice. We asked the question whether these advantages are counteracted by an error related to the intrinsic simplification. We found that axisymmetric DKI is a viable alternative to standard DKI for studies focusing on *D*_⊥_, *D*_‖_ and 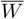 since these parameters could be estimated with few substantially differing voxels (SDV) with respect to their standard DKI counterpart. For all other parameters, i.e., *W*_⊥_, *W*_‖_ and the biophysical parameters, axisymmetric DKI produced results that differed on a larger scale from their standard DKI based counterparts and our study urges caution when planing to use axisymmetric DKI to investigate effects based on those parameters. However, with a median difference of up to 11% for the AxTM and up to 18% for the biophysical parameters, the observed differences might still be in an acceptable range depending on the application. Moreover, the number of SDV can be substantially reduced if an FA threshold is used.

## 6 Ethical Approval Statement

The in-vivo dMRI data used for this study were acquired with the help of a human research participant. The participant provided written informed consent. The local ethics committees at University Medical Center Hamburg-Eppendorf approved the study (PV5141).

## 7 Availability of data and materials

The open source ACID toolbox for SPM contains the estimation methods for standard and axisymmetric DKI. Furthermore, a code repository for generation and analysis of the data used in this study will be made available on Github.

## 9 Authors’ contributions

**Jan Malte Oeschger:** Conceptualization, Data curation, Formal analysis, Investigation, Methodology, Software, Visualization, Writing – original draft, **Karsten Tabelow:** Conceptualization, Methodology, Supervision, Writing – review & editing, **Siawoosh Mohammadi:** Conceptualization, Funding acquisition, Methodology, Project administration, Resources, Supervision, Writing – review

## Glossary

DTI: Diffusion tensor imaging.
DKI: Diffusion kurtosis imaging.
AxTM: Axisymmetric DKI tensor metrics: *D*_‖_, *D*_⊥_, *W*_‖_, *W*_⊥_, and 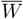.
*D*_‖_: Parallel diffusivity.
*D*_⊥_: Perpendicular diffusivity.
*W*_‖_: Parallel kurtosis.
*W*_⊥_: Perpendicular kurtosis.
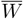: Mean of the kurtosis tensor.

