## Supplementary material for "Investigating apparent differences between standard DKI and axisymmetric DKI and its consequences for biophysical parameter estimates"

June 21, 2023

**1** University Medical Center Hamburg Eppendorf, Institute of Systems Neuroscience, Hamburg, Germany

**2** Weierstrass Institute for Applied Analysis and Stochastics, Berlin, Germany

**3** Max Planck Institute for Human Cognitive and Brain Sciences, Department of Neurophysics, Leipzig, Germany

***** Corresponding author:

| | |
|---|---|
| **Name** | Jan Malte Oeschger |
| **Institute** | University Medical Center Hamburg-Eppendorf, Institute of Systems Neuroscience |
| **Address** | Martinistraße 52, 20246 Hamburg, Germany |
| **E-mail** | |
| **Phone** | +49-40-7410-27301 |

### Supporting Information

#### S1.1 Standard DKI signal representation

For a given diffusion weighting $b$ and diffusion gradient $\vec{g} = (g_1, g_2, g_3)^\intercal$, the noise-free DKI signal can be represented as in[1;2]:

$$\zeta_{b,\vec{g}}(\zeta_0, D, W) = \zeta_0 \exp\left[-bD + \frac{b^2}{6}\left(\frac{\mathrm{Tr}(D)}{3}\right)^2 W\right] \quad \text{[S1.1a]}$$

$$D = \sum_{i,j=1}^{3} g_i g_j D_{ij} \quad \text{[S1.1b]}$$

$$W = \sum_{i,j,k,l=1}^{3} g_i g_j g_k g_l W_{ijkl} \quad \text{[S1.1c]}$$

where $D_{ij}$ are the diffusion tensor entries, $W_{ijkl}$ are the kurtosis tensor entries and $\zeta_0$ is the non-diffusion-weighted signal ($b = 0\frac{\mathrm{s}}{\mathrm{mm}^2}$).

From the tensors D and W, the AxTM can be directly computed: $D_{\|} = \lambda_1$ where $\lambda_1$ is the first eigenvalue of the diffusion tensor D, $D_{\perp} = \frac{\lambda_2+\lambda_3}{2}$. For $W_{\|}$ and $W_{\perp}$, first $K_{\|}$ and $K_{\perp}$ are calculated from the estimated kurtosis tensor elements $W_{ijkl}$ according to Eqs.[3] 31 and 32. $K_{\|}$ and $K_{\perp}$ are then transformed to $W_{\|}$ and $W_{\perp}$ according to Eq. 14 in[4]: $W_{\|} = \frac{K_{\|}D_{\|}^2}{\overline{D}^2}$ and $W_{\perp} = \frac{K_{\perp}D_{\perp}^2}{\overline{D}^2}$ where $\overline{D} = \frac{\lambda_1+\lambda_2+\lambda_3}{3}$ is the mean diffusivity. $\overline{W}$ can be computed according to Eq. 10 from[5]: $\overline{W} = \frac{1}{5}(W_{xxxx} + W_{yyyy} + W_{zzzz} + 2W_{xxyy} + 2W_{xxzz} + 2W_{yyzz})$.

#### S1.2 Axisymmetric DKI

Axisymmetric DKI[4] assumes symmetric diffusion around an axis of symmetry $\vec{c}$ inside an imaging voxel. Mathematically, this assumption leads to axisymmetric diffusion and kurtosis tensors with a drastically reduced number of independent tensor parameters compared to standard DKI (from 15 to 3 parameters for the kurtosis tensor and from 6 to 2 parameters for the diffusion tensor). Apart from the tensors, axisymmetric DKI additionally contains two parameters for the axis of symmetry.

With the axis of symmetry $\vec{c}$ parameterized by the inclination $\theta$ and azimuth $\phi$: $\vec{c} = \begin{pmatrix} \sin\theta\cos\phi \\ \sin\theta\sin\phi \\ \cos\theta \end{pmatrix}$, the diffusion and kurtosis tensors can be determined according to[4]:

$$D = D_{\perp}\mathbf{I} + (D_{\parallel} - D_{\perp})\vec{c}\vec{c}^{T} \qquad \text{[S1.2]}$$

and

$$\begin{aligned} W = &\frac{1}{2}(10W_{\perp} + 5W_{\parallel} - 15\overline{W})\mathbf{P} + W_{\perp}\Lambda + \\ &\frac{3}{2}(5\overline{W} - W_{\parallel} - 4W_{\perp})\mathbf{Q} \end{aligned}$$

where $\Psi = \{D_{\parallel}, D_{\perp}, W_{\parallel}, W_{\perp}, \overline{W}, \zeta_0, \theta, \phi\}$ are the 8 framework's parameters ($\zeta_0$ is the non diffusion-weighted signal) and $\mathbf{I} = \begin{pmatrix} 1 & 0 & 0 \\ 0 & 1 & 0 \\ 0 & 0 & 1 \end{pmatrix}$ is the identity matrix. The tensors $\mathbf{P}, \Lambda$ and $\mathbf{Q}$ can be computed with the Kronecker delta $\delta_{xy}$ and the components of the axis of symmetry $c_x$ ($x, y \in 1, 2, 3$) as: $\mathbf{P}_{ijkl} = c_i c_j c_k c_l$, $\mathbf{Q}_{ijkl} = \frac{1}{6}(c_i c_j \delta_{kl} + c_i c_k \delta_{jl} + c_i c_l \delta_{jk} + c_j c_k \delta_{il} + c_j c_l \delta_{ik} + c_k c_l \delta_{ij})$ and $\Lambda_{ijkl} = \frac{1}{3}(\delta_{ij}\delta_{kl} + \delta_{ik}\delta_{jl} + \delta_{il}\delta_{jk})$[4]. The according noise-free signal $\zeta_{b,\vec{g}}(\Psi)$ can then be computed based upon the axisymmetric tensors[6]:

$$\zeta_{b,\vec{g}}(\Psi) = \zeta_0 \exp\left(-B_{ij}D_{ij} + \frac{1}{6}\overline{D}^2 B_{ij}B_{kl}W_{ijkl}\right) \qquad \text{[S1.3]}$$

where

$$B_{ij}D_{ij} = \mathrm{Tr}(B)D_{\perp} + (D_{\parallel} - D_{\perp})\vec{c}^{T}B\vec{c} \qquad \text{[S1.4]}$$

and

$$B_{ij}B_{kl}W_{ijkl} = \frac{1}{2}(10W_{\perp} + 5W_{\|} - 15\overline{W})(\vec{c}^T B\vec{c})^2 \quad \text{[S1.5]}$$

$$+ \frac{1}{2}(5\overline{W} - W_{\|} - 4W_{\perp})(\vec{c}^T B\vec{c}\,\mathrm{Tr}(B) \quad \text{[S1.6]}$$

$$+ 2\vec{c}^T BB\vec{c}) + \frac{W_{\perp}}{3}(Tr(B)^2 + 2\,\mathrm{Tr}(B \otimes B)) \quad \text{[S1.7]}$$

with

$$B = b\begin{pmatrix} g_x^2 & g_x g_y & g_x g_z \\ g_x g_y & g_y^2 & g_y g_z \\ g_x g_z & g_y g_z & g_z^2 \end{pmatrix}$$

The AxTM can be computed from the standard DKI tensor metrics assuming axial-symmetry, see Section S1.1.

#### S1.3 Derivation of the Relationship between the axial symmetric DKI and the biophysical parameters

The derivation of the relationship between the axial-symmetrical diffusion parameters and the biophysical parameters is based upon the work by[7;8]. Starting point for the derivation are the formulas found in[7] that establish a connection between the axial symmetric kurtosis parameters and the biophysical parameters, based upon the assumption of an axially symmetric fiber orientation distribution function (ODF):

$$M_1 = 3D_0 = fD_a + (1-f)(2D_{e,\perp} + D_{e,\|})$$

$$M_2 = \frac{3}{2}D_2\frac{1}{p_2} = fD_a + (1-f)(D_{e,\|} - D_{e,\perp})$$

$$M_3 = D_2^2 + 5D_0^2(1+\frac{W_0}{3}) = fD_a^2 + (1-f)[5D_{e,\perp}^2 + (D_{e,\|} - D_{e,\perp})^2 + \frac{10}{3}D_{e,\perp}(D_{e,\|} - D_{e,\perp})]$$

$$M_4 = \frac{1}{2}D_2(D_2 + 7D_0)\frac{1}{p_2} + \frac{7}{12}\frac{1}{p_2}W_2D_0^2 = fD_a^2 + (1-f)((D_{e,\|} - D_{e,\perp})^2 + \frac{7}{3}D_{e,\perp}(D_{e,\|} - D_{e,\perp}))$$

$$M_5 = \frac{9}{4}D_2^2 + \frac{35}{24}W_4D_0^2 = p_4(fD_a^2 + (1-f)(D_{e,\|} - D_{e,\perp})^2)$$

M1, M2, M3, M4 and M5 only depend on the biophysical parameter $\kappa$ via the functions $p_2$ and $p_4$ and the axial-symmetrical diffusion parameters $D_\|, D_\perp, W_\|, W_\perp, \overline{W}$:

$$D_0 = \frac{1}{3}(2D_\perp + D_\|) \qquad D_2 = \frac{2}{3}(D_\| - D_\perp)$$

$$W_0 = \overline{W} \qquad W_2 = \frac{1}{7}(3W_\| + 5\overline{W} - 8W_\perp)$$

$$W_4 = \frac{4}{7}(W_\| - 3\overline{W} + 2W_\perp)$$

$$p_2 = \frac{1}{4}(\frac{3}{\sqrt{\kappa}F(\sqrt{\kappa})} - 2 - \frac{3}{\kappa})$$

$$p_4 = \frac{1}{32\kappa^2}(105 + 12\kappa(5+\kappa) + \frac{5\sqrt{\kappa}(2\kappa - 21)}{F(\sqrt{\kappa})})$$

here, F is Dawsons function. A quadratic equation for f can be found (for detailed derivation see[8]):

$$0 = af^2 - (a + c - \frac{40}{3})f + c \qquad \text{[S1.8]}$$

where a and c are:

$$a = (\Delta m)^2 - (\frac{7}{3} + 2d_2)\Delta m + m_2 \qquad \text{[S1.9]}$$

and:

$$c = (\Delta m - 5 - d_2)^2 \tag{S1.10}$$

that depend on $d_2$, $m_2$, $\overline{D}$ and $\Delta m$ which can be computed with the axial-symmetrical diffusion parameters and κ:

$$\overline{D} = \frac{1}{3}(M_1 - M_2) \qquad \Delta m = \frac{M_3}{\overline{D}^2} - \frac{M_4}{\overline{D}^2}$$

$$d_2 := \frac{M_2}{\overline{D}} \qquad m_2 := \frac{M_4}{\overline{D}^2}$$

Eq. (S1.8) has two solutions referred to as "branches", which, in turn, can be computed with $a$ and $c$:

$$f_- = \frac{-\frac{40}{3} + a + c - \sqrt{-4ac + (\frac{40}{3} - a - c)^2}}{2a} \tag{S1.11}$$

$$f_+ = \frac{-\frac{40}{3} + a + c + \sqrt{-4ac + (\frac{40}{3} - a - c)^2}}{2a} \tag{S1.12}$$

The solution for f (either branch "+" or "−") can then be used to compute the diffusivities $D_{e,\perp}$, $D_{e,\|}$ and $D_a$ analytically.

$$D_{e,\perp} = \frac{\overline{D}}{(1-f)} \qquad D_a = (\frac{\Delta m(1-f) - 5 - d_2}{-f})\overline{D}$$

$$D_{e,\|} = (\frac{d_2 - f\frac{D_a}{\overline{D}}}{(1-f)})\overline{D} + D_{e,\perp}$$

However, at this point κ is still unknown and needed to estimate $p_2$ and $p_4$. All the biophysical diffusion parameters and f can now be expressed in terms of κ and the axial-symmetrical diffusion parameters. This is used to define an objective function where κ is the only unknown parameter, since the axial-symmetrical diffusion parameters have previously been estimated:

$$0 = [p_4(fD_a^2 + (1-f)(D_{e,\parallel} - D_{e,\perp})^2)] - M_5 \quad \text{[S1.13]}$$

This objective function is then minimized to find κ with which first f and then the biophysical diffusivites can be found as described.

#### S1.4 Comparison of fit of log of signals with NLLS fit

It was analytically shown[9] that standard DKI and axisymmetric DKI produce the same results under two conditions. The first condition was that the log of the signals are being fitted. To rule out the possibility that the observed differences as measured by the A-PE between standard DKI and axisymmetric DKI in this study are caused by fitting the non-linear signals, a log-of-signals fit was implemented for both standard DKI and axisymmetric DKI and used to fit the same dMRI data described in Section 2.1 (main document). Figure S1 shows the difference between standard DKI and axisymmetric DKI when fitting the log-of-signals (red histogram) versus the differences when using the NLLS fit implementation (blue histogram) on top and the scatter density plots including between the results obtained with both methods at the bottom.

Fitting the log of the signals still went along with substantial differences between both models that in some cases, e.g. $W_\perp$, showed a close relation with the NLLS fit results. It can therefore be ruled out that the observed differences in the main study are purely caused by not fitting the log of the signals.

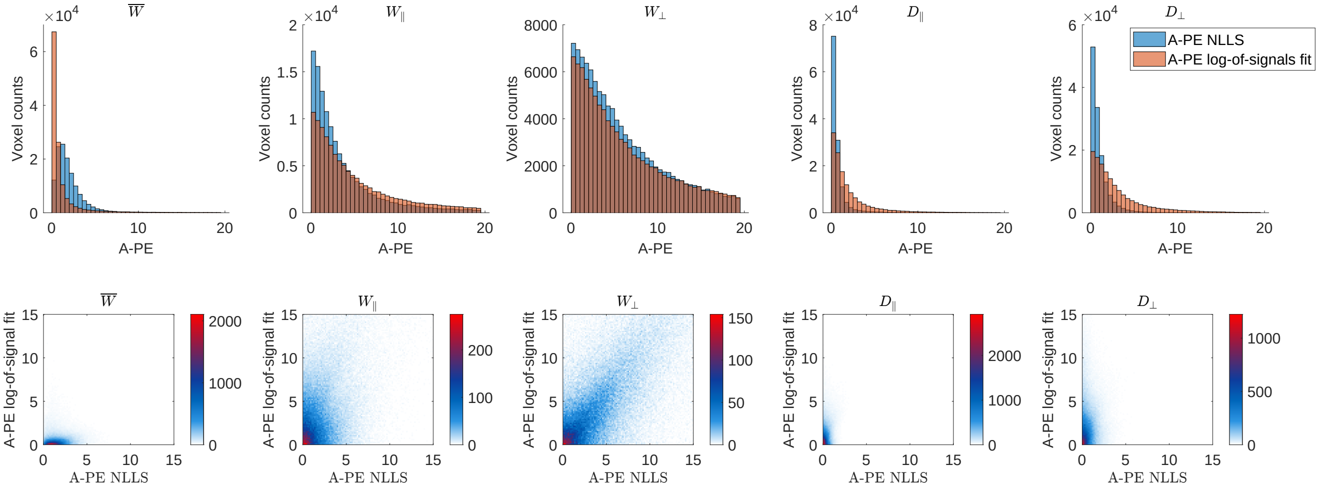


Figure S1: A-PE estimated based on a log-of-signals fit and the non-linear least sqaures (NLLS) fit used for this study. Top: histograms of A-PE distributions when fitting the log-of-signals (red histogram) versus the NLLS fit implementation (blue histogram). Bottom: Scatter density plots of the A-PE estimated with a log-of-signals fit versus the NLLS fit.

#### S1.5 Histograms of A-PE for the AxTM and biophysical parameters

Because the main document cites the summary measures number of substantially differing voxels (SDV) and median bias in the population of SDV for brevity, Figure S2 shows the actual distributions of A-PE for the AxTM and biophysical parameters.

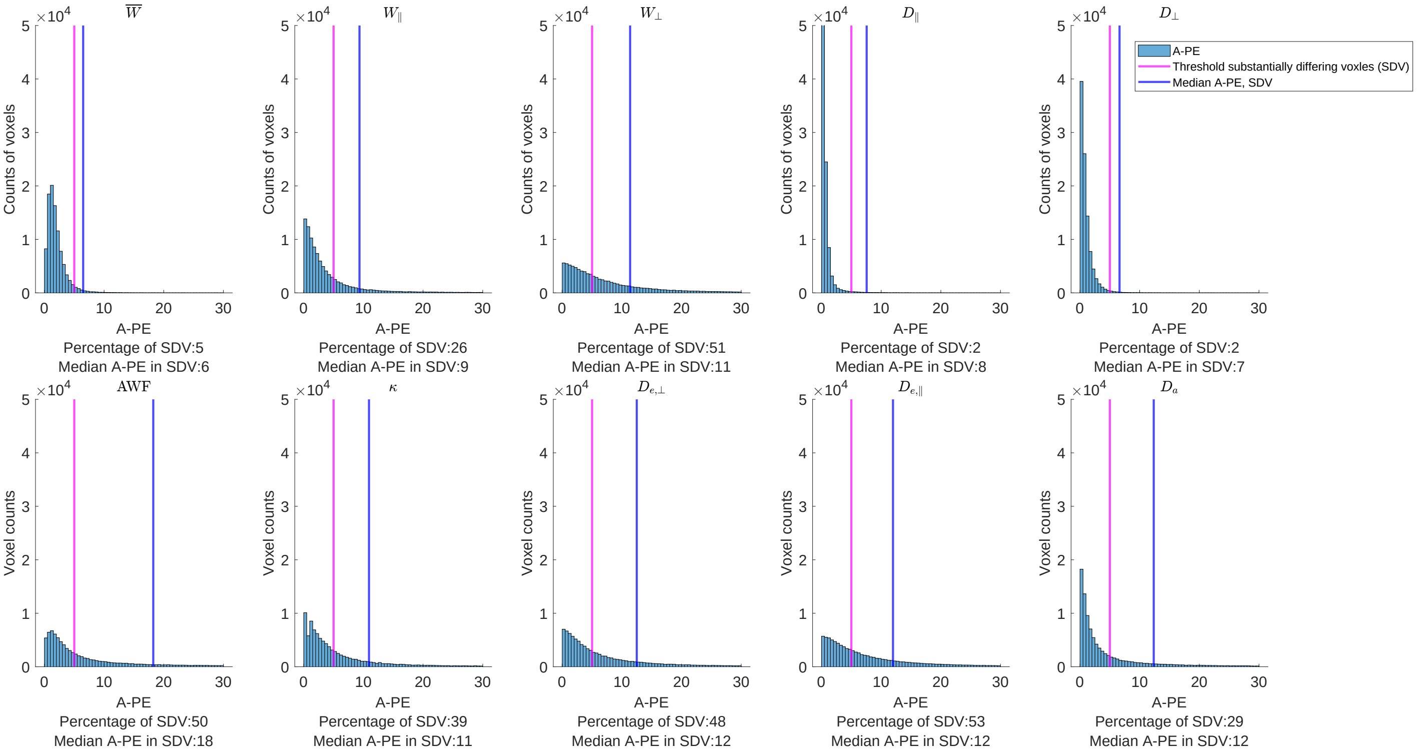


Figure S2: Histograms of underlying A-PE distribution for the AxTM and biophysical parameters. The pink vertical lines indicate the threshold after which axisymmetric DKI parameter estimation results were considered "substantially differing" (A-PE $\geq$ 5%), the blue vertical lines indicate the median difference in the population of substantially differing voxels. The x-axis was confined to [0 30].

##### S1.6 Inter-dependence of A-PE and difference in main fiber orientation

To establish whether the difference between the main fiber orientation found by both DKI models is causing the difference between fit results of both models, Figure S3 documents the scatter density plots of A-PE and angle $\phi$. A-PE and angle $\phi$ showed the biggest inter-dependency for $W_{\parallel}$ and $W_{\perp}$ allowing to establish at least a partial causality in this case.

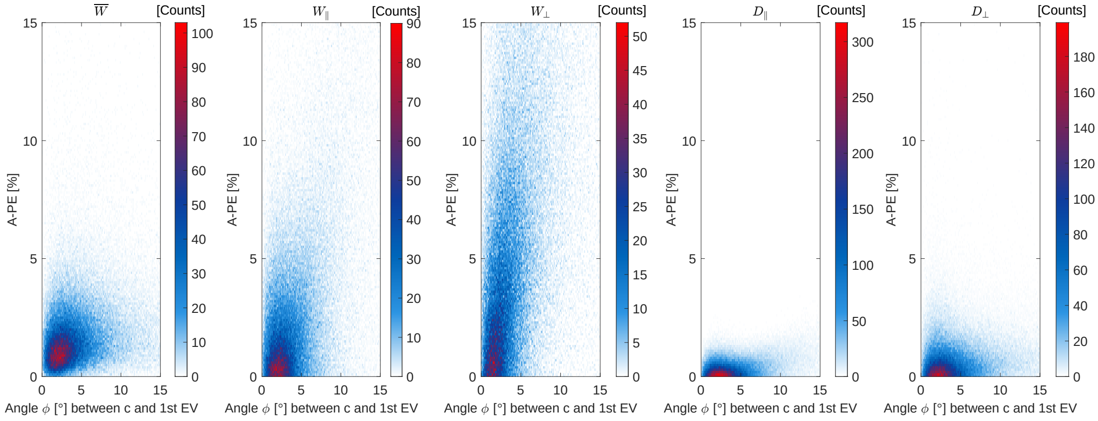


Figure S3: Scatter density plots between A-PE and angle $\phi$ (see Section 2.3, main document), computed for voxels in the white matter mask.

[3] A. Tabesh, J. H. Jensen, B. A. Ardekani, and J. A. Helpern, "Estimation of tensors and tensor-

derived measures in diffusional kurtosis imaging," *Magnetic Resonance in Medicine*, vol. 65, no. 3, pp. 823–836, 2011.

[4] B. Hansen, N. Shemesh, and S. N. Jespersen, "Fast imaging of mean, axial and radial diffusion kurtosis," *NeuroImage*, vol. 142, pp. 381–393, Nov. 2016.

[5] B. Hansen, T. E. Lund, R. Sangill, and S. N. Jespersen, "Experimentally and computationally fast method for estimation of a mean kurtosis," *Magnetic Resonance in Medicine*, vol. 69, no. 6, pp. 1754–1760, 2013.

[6] B. Hansen, A. R. Khan, N. Shemesh, T. E. Lund, R. Sangill, S. F. Eskildsen, L. Østergaard, and S. N. Jespersen, "White matter biomarkers from fast protocols using axially symmetric diffusion kurtosis imaging," *NMR in Biomedicine*, vol. 30, no. 9, p. e3741, 2017.

[7] S. N. Jespersen, J. L. Olesen, B. Hansen, and N. Shemesh, "Diffusion time dependence of microstructural parameters in fixed spinal cord," *NeuroImage*, vol. 182, pp. 329–342, 2018.

[8] "Rotationally-invariant mapping of scalar and orientational metrics of neuronal microstructure with diffusion MRI," *NeuroImage*.

[9] S. Nørhøj Jespersen, "White matter biomarkers from diffusion MRI," *Journal of Magnetic Resonance*, vol. 291, pp. 127–140, June 2018.
